## Supplementary figures and images for "ScRNA-seq Expression of *IFI27* and *APOC2* Identifies Four Alveolar Macrophage Superclusters in Healthy BALF"

### Figure S1

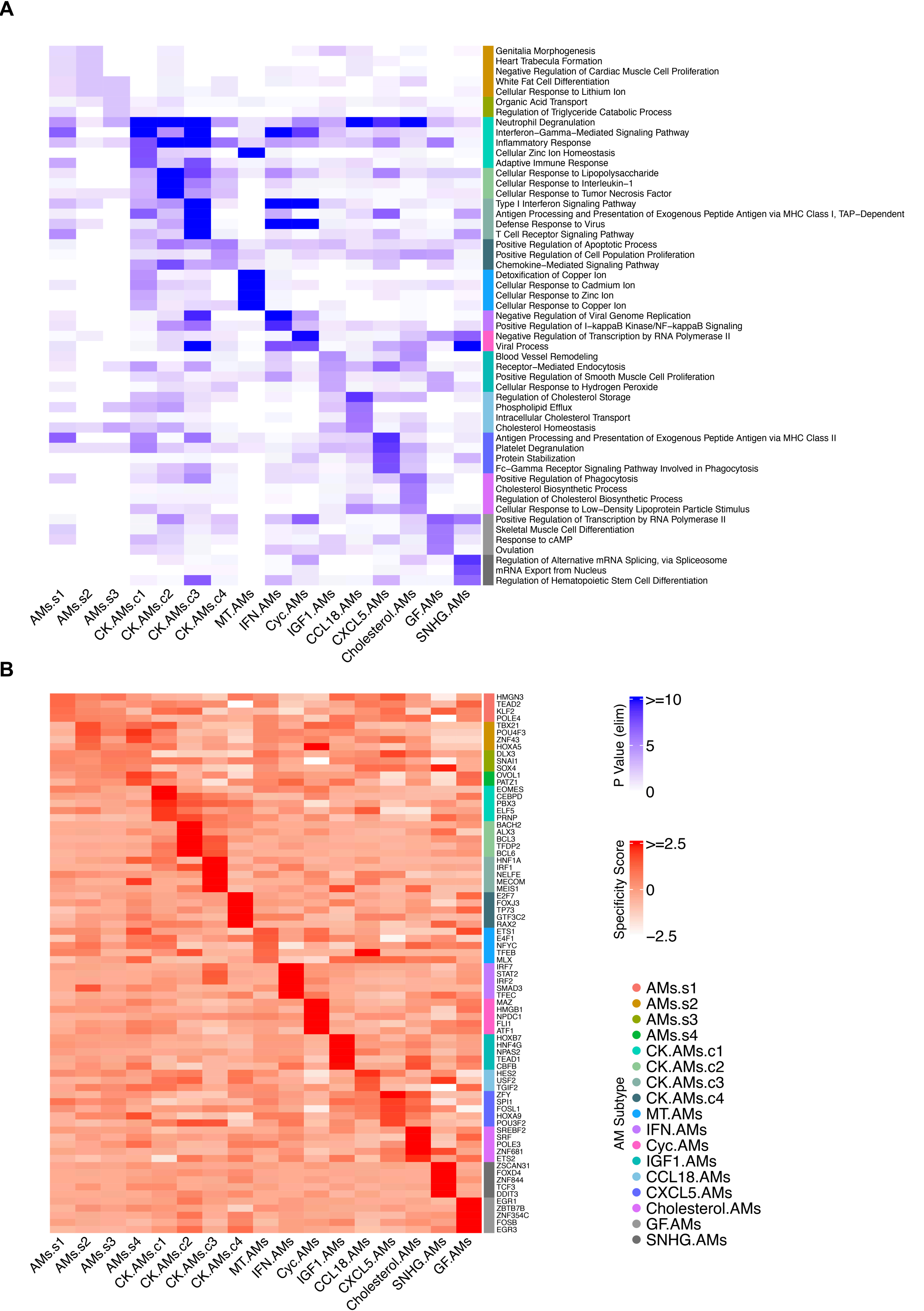

### Figure S2

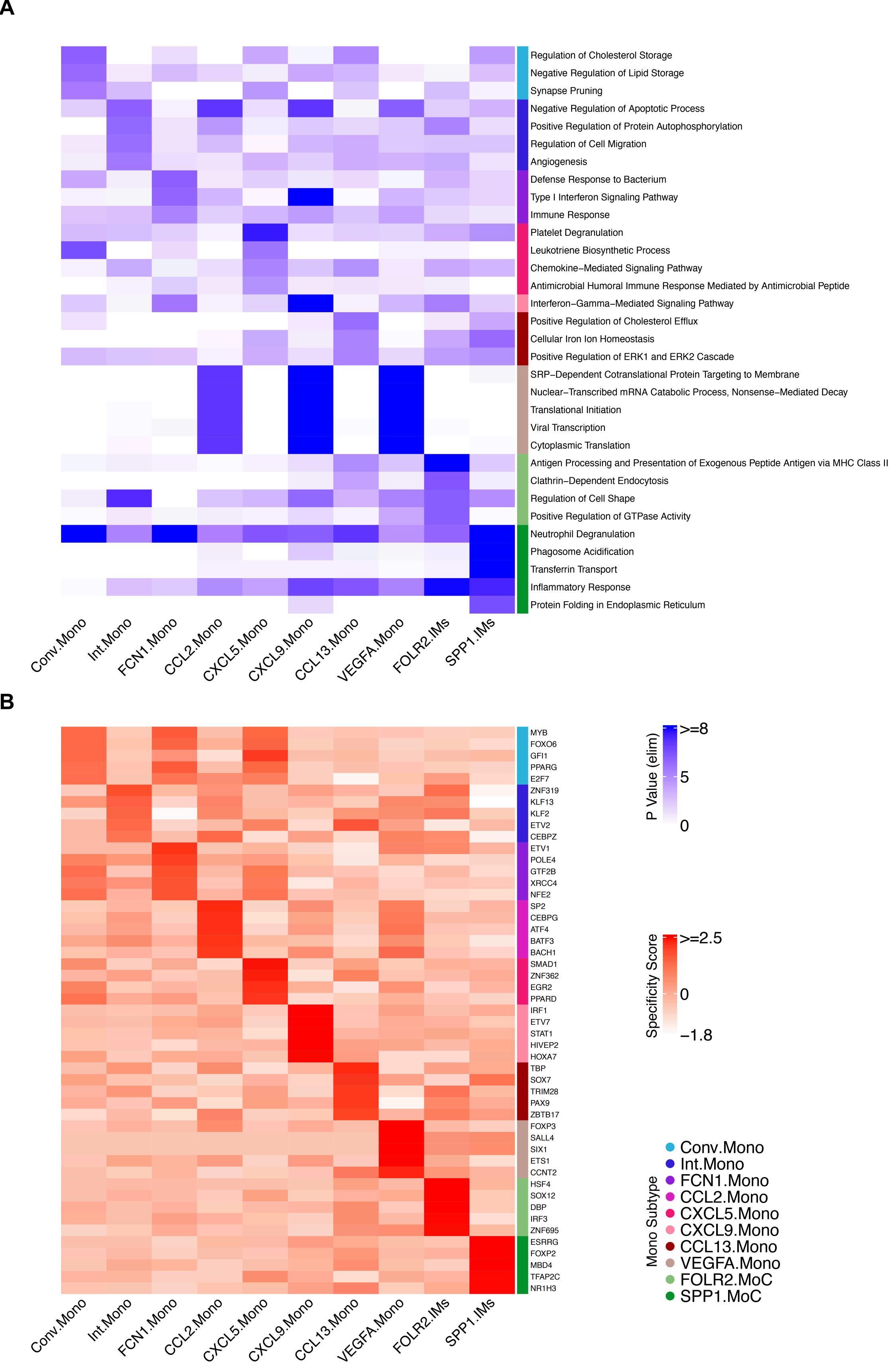

### Figure S3

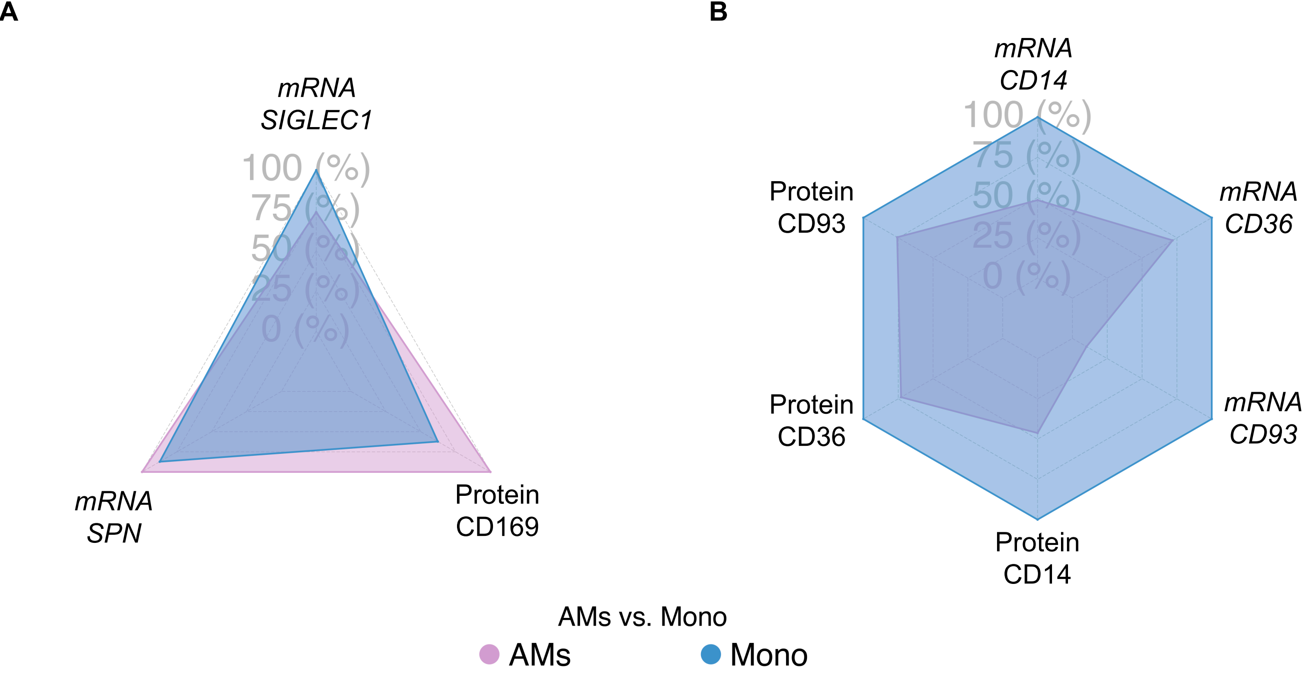

### Figure S4

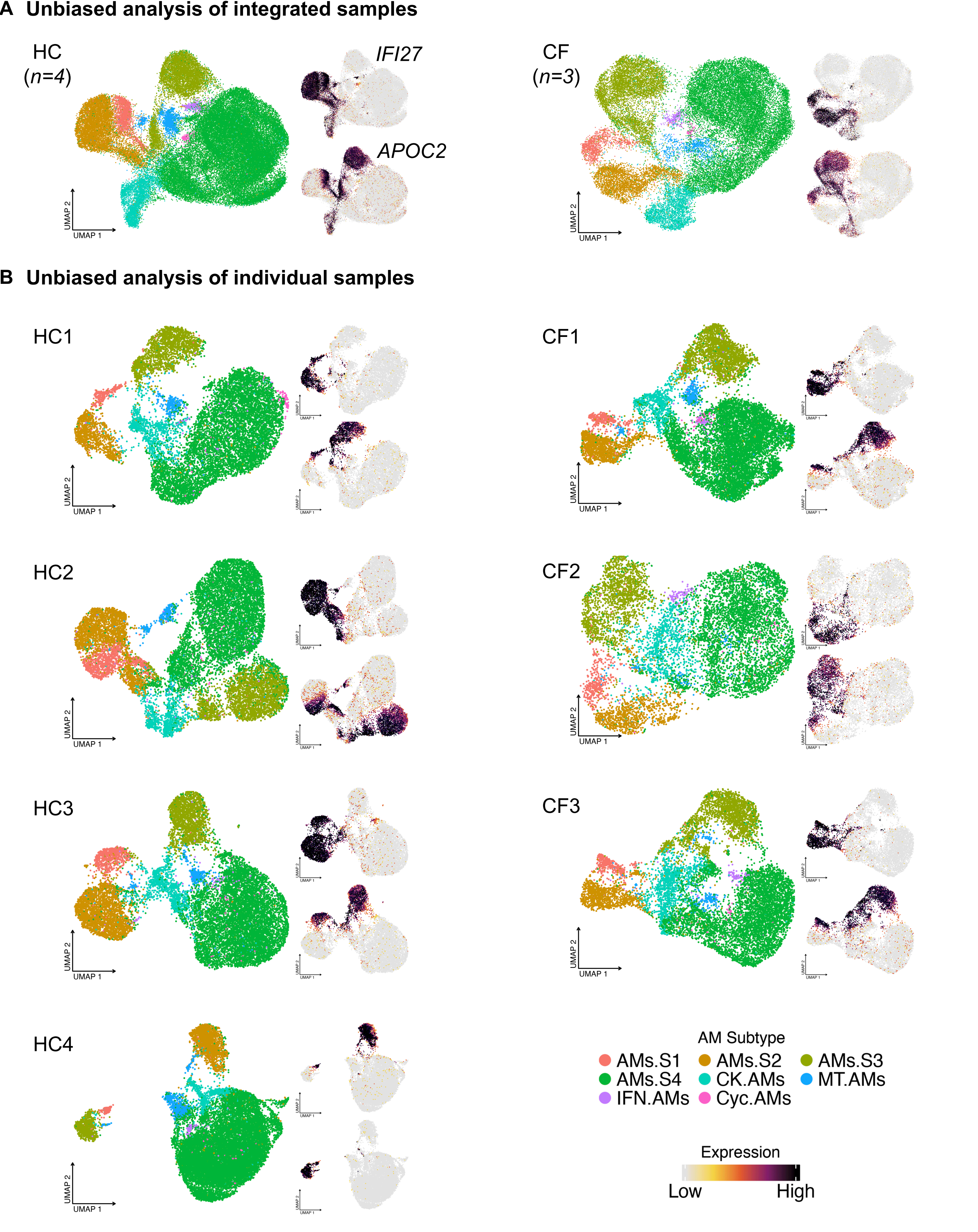

### Figure S5

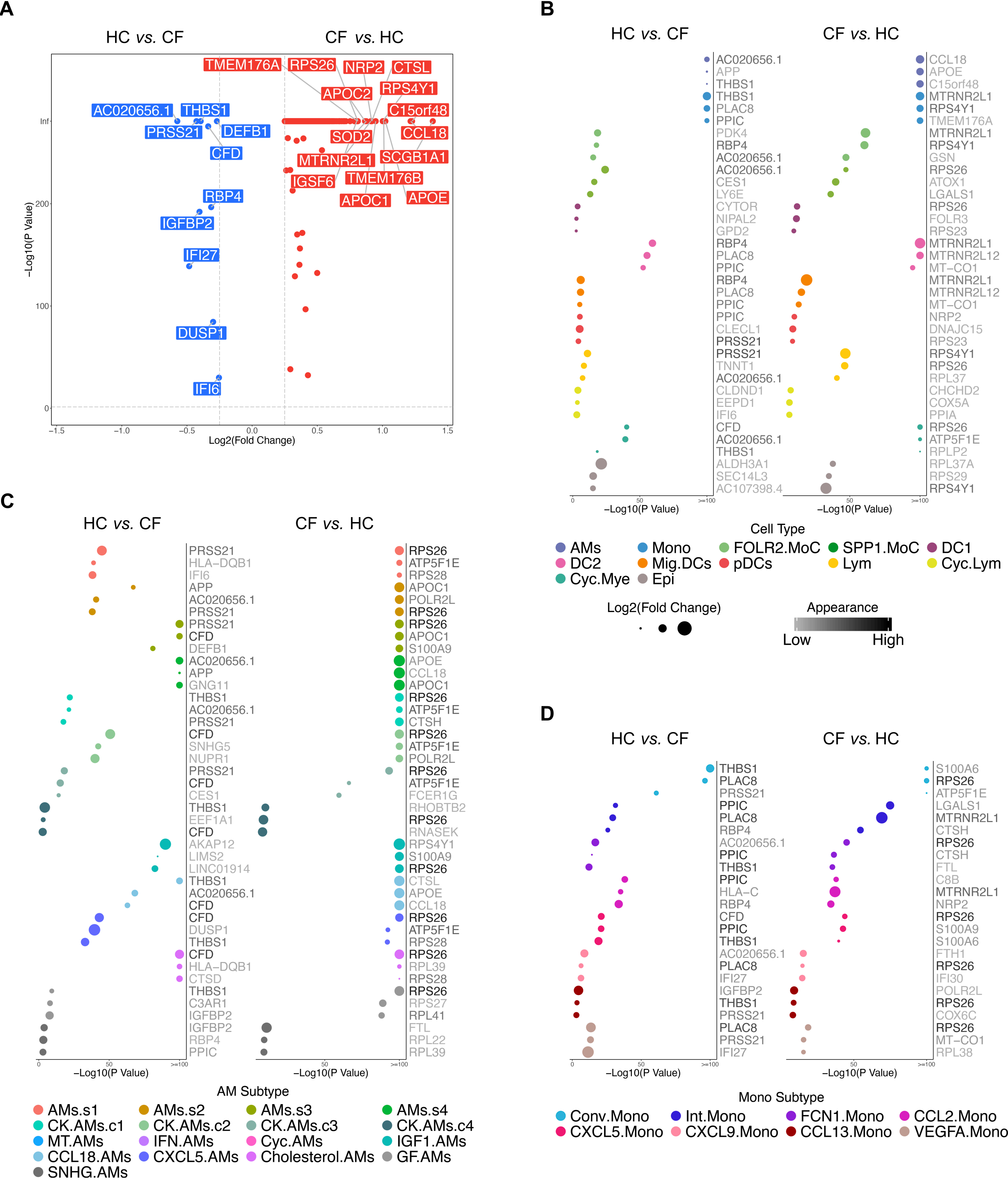
