## Supplementary material for "ScRNA-seq Expression of *IFI27* and *APOC2* Identifies Four Alveolar Macrophage Superclusters in Healthy BALF": Figure S6

A

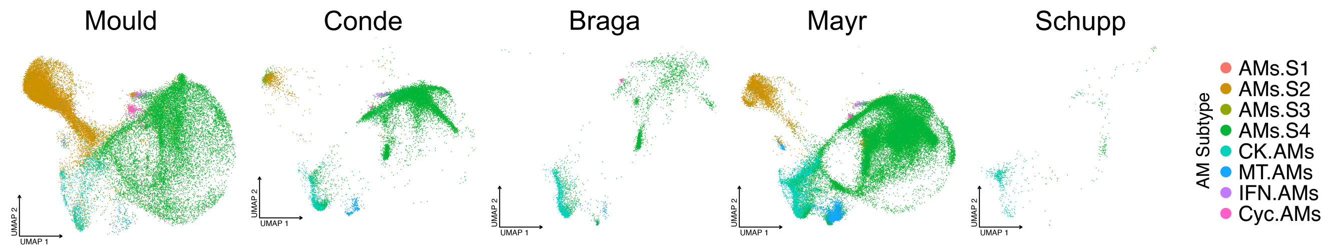


B

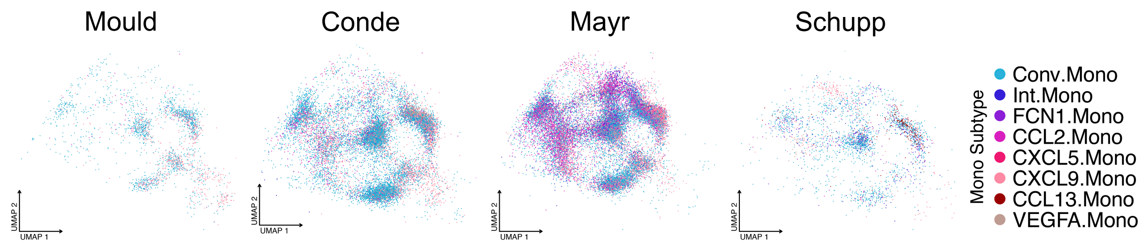


C

| Study | Tissue | Health Status | Cell No. | Reference |
|---|---|---|---|---|
| Li | BAL | HC/CF | 9370 | |
| Mould | BAL | HC | 2903 | PMID: 33079572 |
| Conde | Deceased lung piece | HC | 24267 | PMID: 35549406 |
| Mayr | Fresh lung piece | HC/Endstage Fibrosis | 14603 | PMID: 33650774 |
| Schupp | Sputum | HC/CF | 2391 | PMID: 32603604 |

D

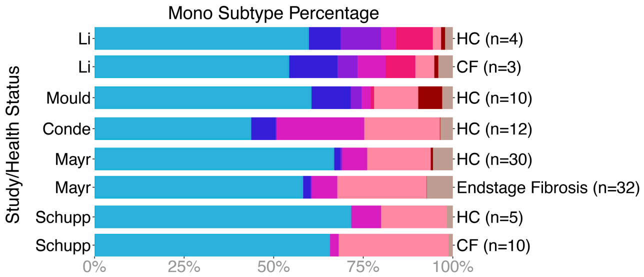


E

F

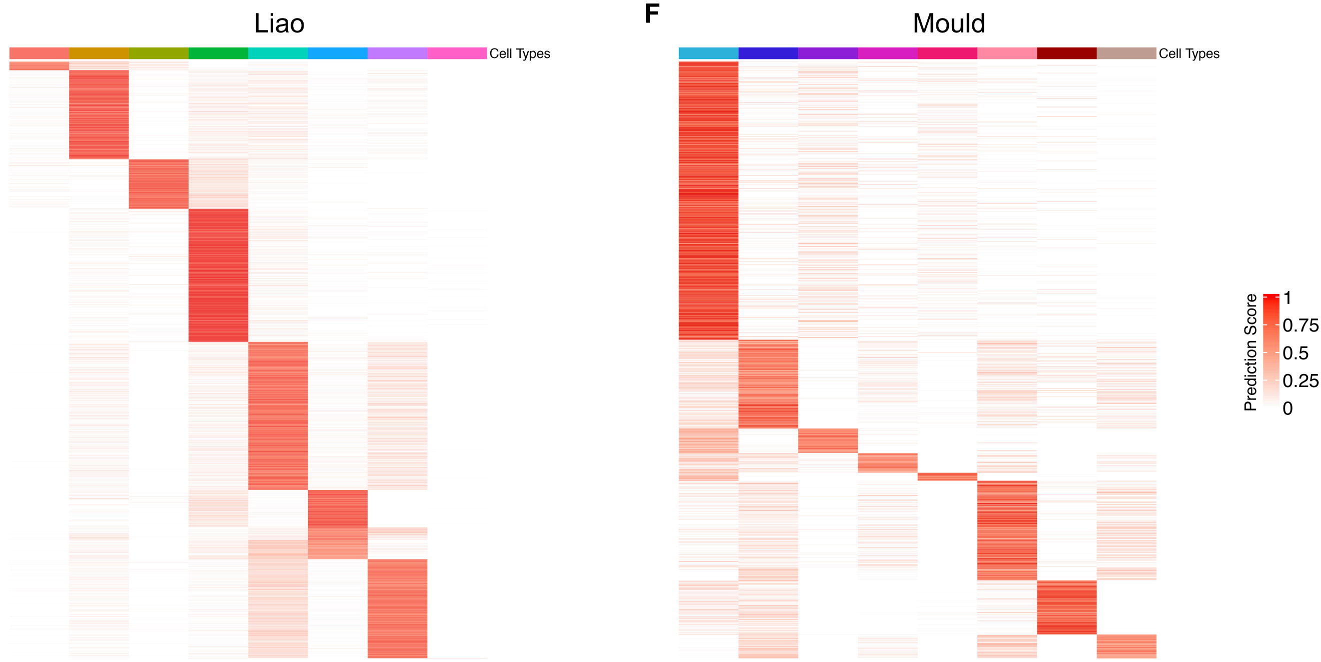


Fig. S6
